# Toward transparent taxonomy: an interactive web-tool for evaluating competing taxonomic arrangements

**DOI:** 10.1101/2023.06.20.545819

**Authors:** Oksana V. Vernygora, Felix A.H. Sperling, Julian R. Dupuis

## Abstract

1. Informative and consistent taxonomy above the species level is essential to communication about evolution, biodiversity, and conservation, and yet the practice of taxonomy is considered opaque and subjective by non-taxonomist scientists and the public alike. While various proposals have tried to make the basis for ranking and inclusiveness of taxa more transparent and objective, widespread adoption of these ideas has lagged.

2. Here, we present TaxonomR, an interactive online decision-support tool to evaluate alternative taxonomic classifications. This tool implements an approach that quantifies the criteria commonly used in taxonomic treatments and allows the user to interactively manipulate weightings for different criteria to compare scores for taxonomic groupings under those weights.

3. We use the butterfly taxon *Argynnis* to demonstrate how different weightings applied to common taxonomic criteria result in fundamentally different genus-level classifications that are predominantly used in different continents and geographic regions. These differences are objectively compared and quantified using TaxonomR to evaluate the kinds of criteria that have been emphasized in earlier classifications, and the nature of the support for current alternative taxonomic arrangements.

4. The main role of TaxonomR is to make taxonomic decisions transparent via an explicit prioritization scheme. TaxonomR is not a prescriptive application. Rather, it aims to be a tool for facilitating our understanding of alternative taxonomic classifications that can, in turn, potentially support global harmony in biodiversity assessments through evidence-based discussion and community-wide resolution of historically entrenched taxonomic tensions.

## Introduction

Modern taxonomy is a fundamental biological discipline that governs the names, descriptions, and classification of the natural world (Raven et al., 1971, Wheeler, 2008, Wheeler et al., 2004, Wilson, 2004). Its principles and practice were first set in motion by Carl Linnaeus (1735) almost three centuries ago and now permeate communication, conservation, and collaboration across a global network. Modern taxonomy extends far beyond mere “stamp collecting” to encompass a wide range of knowledge about organisms including their evolutionary history, biology, distribution, and ecology (Schram, 2004, Vences et al., 2013, Wheeler et al., 2004, Wilson, 2004). Taxonomy lays the foundation for all other fields of biology by organizing the natural world into discrete hierarchically arranged units of living organisms that are easy to identify, manage, and protect. And yet, despite its indispensable role in biodiversity information management, it has been deemed a vague, poorly understood, archaic and even anarchic science (Agnarsson and Kuntner, 2007, Garnett and Christidis, 2017).

Taxonomy’s stigma of opacity and inconsistency is partially due to a lack of understanding of the conventions under which the field operates above the species level. Owing to intrinsic differences among the diverse forms of life, as well as our interactions with them, there is no single way that taxonomic groups have been delineated uniformly across all organisms and all classification purposes. Therefore, in practice, taxonomic ranking decisions are a product of the evaluation and weighting of a number of criteria related to the inferred relatedness, appearance, and life history characteristics of a given group of organisms. Mayr and Ashlock (1991) described seven commonly used criteria that they referred to as a group’s distinctness and uniqueness (relative size of gap), degree of difference (overall divergence between groups), evolutionary role, grade characteristics (shared adaptive zone or organizational level), size of a group (number of taxa included in that group), equivalence of ranking in related taxa, and taxonomic stability. However, they provided no formal framework for weighting these criteria in delineations of taxonomic groups, other than a “balanced consideration” in which the respective importance of criteria can differ from case to case (Mayr and Ashlock, 1991). More recently, Vences et al. (2013) have provided an extensive list of criteria for naming taxa along with a general workflow aimed to reduce subjectivity in taxonomic practice. Nonetheless, the degree to which some criteria are prioritized over others remains highly subjective, varying not only across research areas but also between individual investigators. Such subjectivity in delimitation and decisions about ranking has led to taxonomic inconsistencies at multiple levels that have become more globally visible in recent literature (Avise and Liu, 2011, Braby et al., 2012, Zhang et al., 2020). Crucially, the lack of clearly defined operational guidelines makes it difficult, if not impossible, to replicate previous decisions or reconcile discrepancies among alternative ranking schemes. Such vagueness and inconsistency challenge taxonomy as valid science in light of a currently growing demand for improved reproducibility and replicability (Allison et al., 2018, Baker, 2016, Shiffrin et al., 2018).

The need for more rigorous governance of taxonomy has been raised in recent literature urging strict adherence to the rules of existing regulatory organizations (Garnett and Christidis, 2017, Garnett and Christidis, 2018, Thomson et al., 2018). However, these bodies, including the International Commission on Zoological Nomenclature (ICZN), the International Association for Plant Taxonomy (IAPT), International Committee on Systematics of Prokaryotes (ICSP), and International Committee on Taxonomy of Viruses (ICTV), only regulate the formation and consistent application of names, and are silent on how to determine the boundaries of taxa, except for the role of the holotype specimen that must be included in a taxon for a name to apply. Further, except for widespread acceptance of the principle that higher level taxa must be monophyletic to be formally recognized, studies rarely assess the consistency and reasoning underlying the delimitation of higher-level taxa (i.e., whether and at what rank to formally name a branch on the tree of life) and there is currently no tool to objectively evaluate and compare alternate taxonomic classifications for groups of species.

A potential solution for taxonomy may be borrowed from other areas of research and industry where decision making processes are facilitated and made transparent through the use of decision support tools (Bhargava et al., 2007, Power, 2007). These applications record explicit user-defined weights and preferred options, combine them in some form or fashion, and output a decision based on the overall prioritization scheme. This general approach has been successfully used in conservation biology, clinical research, industry, and other fields (e.g., Bessette et al., 2019, Copp et al., 2021, Hemming et al., 2022, Morales-Torres et al., 2016, Wilby et al., 2002). The major benefit of using such decision support tools lies in making the decision process transparent and trackable without enforcing the rigid constraints of a single solution to all cases. In a field as diverse as taxonomy, a more objective and quantitative method is urgently needed to support taxonomic decisions and make them more transparent and user-friendly. Establishing such a decision support tool would also help to alleviate the scepticism toward taxonomy that is expressed among both the broader scientific community and the general public (Padial et al., 2010).

Here, we propose an approach implemented with an interactive decision support tool, TaxonomR, that helps to visualize and assess taxonomic decisions based on several criteria traditionally used in taxonomy. This approach aims to facilitate comparability among alternate classifications based on the weights of criteria used to make taxonomic decisions. Conversely, this tool can allow an assessment of the relative weights that must be applied to criteria to allow conformance with different taxonomic arrangements. Our online application can also be used as an effective teaching tool to introduce students to the decision-making process in taxonomy and to provide a quantitative measure of taxonomic classifications for improved reproducibility and replicability of taxonomic and systematic studies. TaxonomR is built in an R environment, publicly available as a shiny app, and is both flexible and interactive for various research- and teaching-based foci.

### Case study: the greater fritillary butterflies

To demonstrate an application of TaxonomR for research, we use it to explore the contentious taxonomy of about 40 species of charismatic greater fritillary butterflies that have been variably classified for over 250 years but are now commonly either placed into a single genus, *Argynnis* Fabricius, 1807 (e.g., Pelham, 2023, Warren et al., 2023, Wikipedia, 2023b, Zhang et al., 2020) or into three major groups ranked as genera: *Argynnis*, *Speyeria* Scudder, 1872, and *Fabriciana* Reuss, 1920 (e.g., De Moya et al., 2017, Savela, 2023, Wahlberg, 2020, Wikipedia, 2023a). In addition, several taxa within *Argynnis* sensu lato (s.l.) that were even more narrowly circumscribed have been treated as distinct genera in many global lists and regional works, including *Argyreus* Scopoli 1777, *Argyronome* Hübner 1818, *Childrenia* Hemming 1943, *Damora* Nordmann 1851, *Mesoacidalia* Reuss 1926, *Nephargynnis* Shirosu & Saigusa 1973, and *Pandoriana* Warren 1942 (e.g., Higgins, 1975, Smart, 1975).

Separation of multiple Eurasian genera within *Argynnis* s.l. was started by Reuss (1926) and supported by Warren (1955, 1944) and Higgins (1975). This resulted in placements of species into numerous different genera until the 1980’s in Europe (Higgins and Riley, 1983) and the 2000’s in Asia (Gorbunov and Kosterin, 2007, Kehimkar, 2008, Shou et al., 2006). Following work by Dos Passos and Grey (1945) in North America, *Speyeria* was usually treated as a distinct genus until revision of a widely followed list by Pelham (2023). The perceived genitalic distinctiveness of each of the major and minor lineages within *Argynnis* s.l. served as the primary justification for recognizing them at the genus rank within Europe and Asia (Gorbunov, 2001, Warren, 1944). One additional argument for splitting *Speyeria* from *Argynnis* was the North American geographic distribution of *Speyeria*, as compared to the Eurasian distribution of the rest of *Argynnis* s.l. (Dos Passos and Grey, 1945). However, recent phylogenetic studies showing a relatively low degree of divergence between these lineages, as well as a lack of consistent reciprocal monophyly among the members of some groups, have revived the idea that only a single genus, *Argynnis*, should be recognized, with *Speyeria* and *Fabriciana* at best being treated as subgenera (e.g., Simonsen et al., 2006, Simonsen, 2006, Zhang et al., 2020).

Here, we use a recent phylogeny (De Moya et al., 2017) of *Argynnis* s.l. (including *Speyeria* and *Fabriciana*) to show how different prioritizing schemes affect taxonomic ranking decisions. We explore multiple approaches to taxonomic classifications, including those that stress phylogenetic or more traditional frameworks, stability in name usage, and those focussed on molecular dating of phylogenies. Explicit weighting of key decision-making criteria illustrates how alternative taxonomic classifications are derived and justified despite using the same set of underlying input information (e.g., phylogeny, biogeography, ecological traits). We compare prioritization schemes under alternate classifications of *Argynnis* and discuss their broader implications for the future of taxonomic research.

## Methods

TaxonomR is an interactive application designed to assess taxonomic ranking decisions based on criteria commonly used in taxonomic practice. This interactive tool (Figure 1) allows users to set weights for each criterion, making the prioritization scheme explicit, transparent, and comparable across different studies. The program is developed using the Shiny package for R programming language and is available as an online application (https://oksanav.shinyapps.io/TaxonomR/) and as a downloadable stand-alone package along with the source code at https://github.com/OksanaVe/TaxonomR.

**Figure 1.**
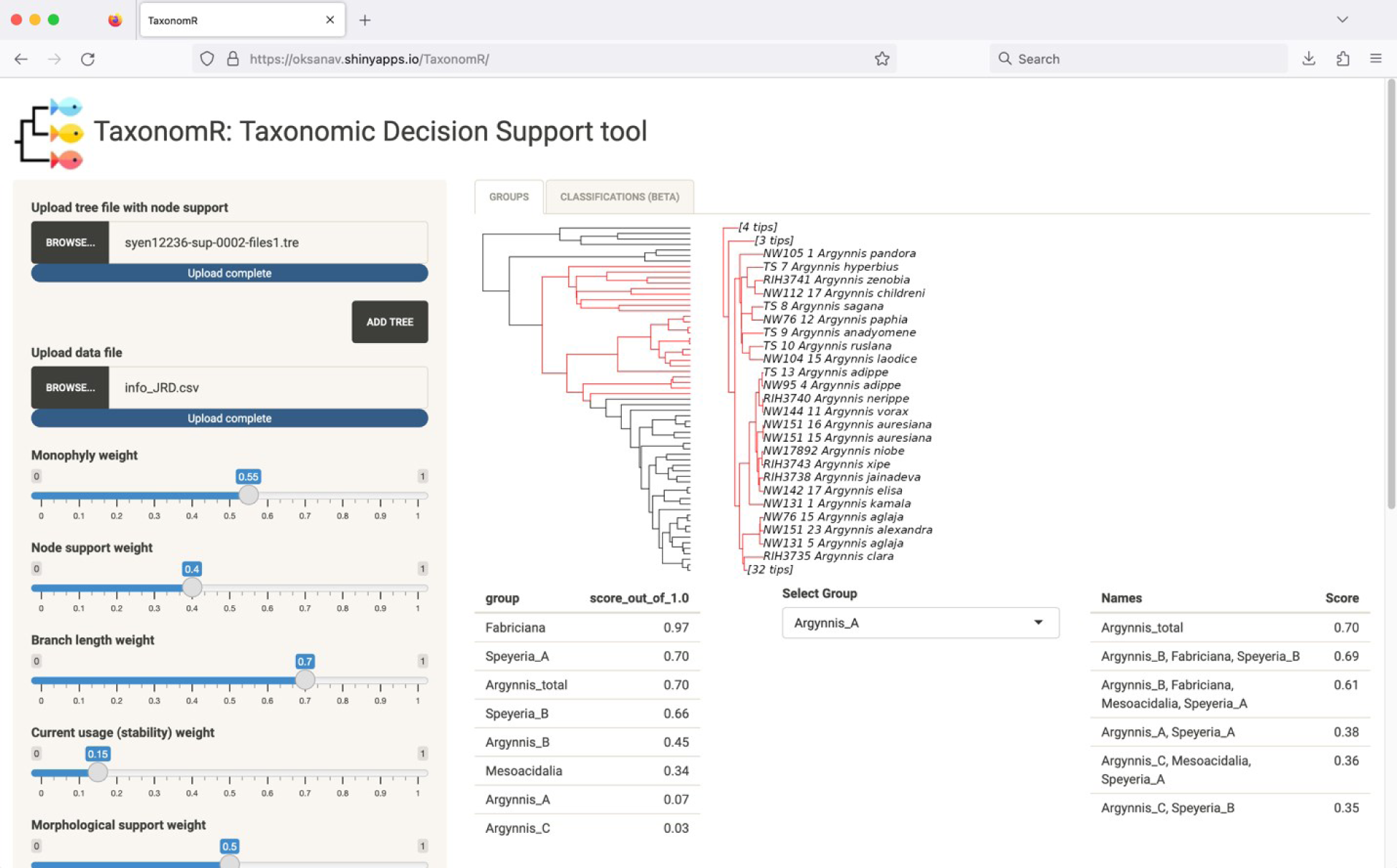
TaxonomR shiny app running in a web browser. Left panel shows file upload and weight sliders, and right panel shows uploaded phylogeny, subtree for a selected group, and scores for individual groups and taxonomic classifications.

### Input data types

TaxonomR requires two input files: a tree file and a CSV data file. Examples of these files are found as supplementary data as well as on the project’s GitHub page.

The **tree file** can be in nexus or newick format and should contain a single tree with branch lengths and support values (if two types of support values are present, TaxonomR will read the first value listed). Optional annotations such as node ages can also be included as in standard output tree files from MrBayes, BEAST, IQ-TREE and other commonly used phylogenetic inference programs (and are required to consider clade age in score calculation). Up to two tree files can be uploaded into the program if support values and divergence times are estimated separately. In this case, both files should include the same topology with the first tree file containing node support values and the second file containing a time-calibrated tree. It is important to note that TaxonomR is not software for phylogenetic inference and does not reconstruct topologies from data files; instead, it requires an input tree file produced by an external program for phylogenetic inference.

(2) The **data file** is a CSV table that contains information about the groupings of taxa. For each proposed taxonomic group (e.g., genus, family, order), the following information should be provided in a single row under a column header with the following corresponding columns (note that all entries in the data file are case-sensitive):

–*Group*: name of a taxonomic group (e.g., genus name);

–*Taxa*: list of terminal taxa included in a group (e.g., list of species included in the genus).

Taxon names should be spelled exactly as they appear in the tree file;

–*Usage*: this field includes information on the current usage of the proposed taxonomic group – it is scored as “new” for a newly proposed taxonomic group name, “new_use” for a previously used taxonomic name but with a modified taxonomic composition, “minor_use” for an existing but rarely used taxonomic name, and “major_use” when using a previously established and commonly used taxonomic name;

– *Morph*: this field indicates presence or absence of distinct morphological traits characterizing a proposed group. For operational convenience, this field is scored on a discrete scale with possible states defined as either “none” for the absence of distinguishing phenotypic traits (e.g., cryptic taxa that are genetically differentiated but are not morphologically distinguishable), “plastic”, if morphological features that characterize a taxon are plastic continuous traits such as colouration or highly plastic meristic and morphometric characters prone to homoplasy (e.g., number of gill rakers and body ratios in fish), “combo” for a unique combination of non-unique phenotypic traits (e.g., trout-perches of the genus *Percopsis* are characterized by the presence of an adipose fin, ctenoid scales, and fin spines; individually, these features are not unique to the group but together form a unique diagnostic combination), and “synap” for the presence of one or more unambiguous uniquely derived morphological traits that characterize a group.

– *Age*: this value specifies the estimated age of the most recent common ancestor of all terminal taxa included in a group and is calculated as the age of the oldest divergence within the proposed taxonomic group. A single value is entered in this column, which initiates an additional slider bar in the program interface where standard deviations can be indicated. The age value should be in the same units as the nodal age values in the tree file.

– *Distribution*: geographical range of the group. Distribution can be specified as a list of biogeographical realms, countries, regions, or areas in which members of the group occur. Range units should not include extensively overlapping or mutually inclusive regions (e.g., New World and South America); they can include areas of any scale, from provincial regions to countries and entire continents.

– *Ecology*: ecological trait indicative of the group’s adaptive zone. It can include information on the diet, foraging habit, reproductive strategy, type of locomotion, nesting and migratory behaviour, host association, or other traits that may be important in defining taxonomic groups. This criterion must be scored as discrete traits and input into the information spreadsheet using short and descriptive keywords without any spaces, e.g., type of habitat can be scored as “woodlands”, “tall_grasslands”, “tropical_rainforest”, “tundra”, etc.

– *Miscellaneous custom traits:* current implementation of TaxonomR allows users to specify up to two additional traits in the “Misc_trait” columns of the data file. While the choice of traits is determined entirely by the user, it should be clearly defined in the study. Scoring and formatting of these traits are similar to that of “Ecology”.

Data fields for any column except Group and Taxa can be left blank if the information for a given criterion is not available or is not considered important.

### Taxon recognition criteria

There are eight criteria for taxon recognition in the current implementation of TaxonomR: monophyly, clade support (node support), distinctiveness (via branch length), current usage (stability), phenotypic recognizability (morphology), clade age, biogeography (distribution), and ecological similarity. These criteria for taxon delimitation and ranking have previously been designated as key criteria for making taxonomic ranking decisions (Mayr and Ashlock, 1991, Vences et al., 2013); however, no formal quantitative framework has been proposed for making arbitrary weighting schemes more explicit. The criteria used in the decision process are not independent of each other and can overlap to various degrees. The information necessary to score these criteria is extracted from the input tree and/or data files.

– *Monophyly* is a central criterion in both evolutionary and cladistic taxonomy and relies directly on a phylogenetic tree. By this criterion, each taxon should represent a monophyletic group. Vences et al. (2013) considered monophyly to be the only strict and necessary taxon naming criterion, emphasising that only monophyletic clades represent natural groups. While this criterion is of primary, if not exclusive, importance for cladistic classifications, its role may be secondary for some special-purpose classifications used for public engagement, folk taxonomy, conservation, and environmental management (Moritz, 1994). Monophyly also carries with it varying degrees of confidence, which we recognize as clade support, and so the strictness of this criterion is potentially reduced for groupings with low support.

– *Node support* is represented by bootstrap or posterior probability values associated with the nodes supporting clades in a phylogenetic tree. Because clades are monophyletic groups by definition, the algorithm only considers node support for monophyletic groups.

– *Branch length* is used as a proxy of distinctiveness of a group and is determined by the “evolutionary distance” between taxonomic groups. In practice, this criterion is calculated as a ratio between length of the longest internal branch within a clade and that of the stem branch leading to the group. The lower the ratio, the higher the group’s distinctiveness is.

– *Current usage (stability)* of a taxonomic classification is an important criterion that directly impacts the usefulness of taxon names and classification as a whole. Many international nomenclatural commissions specify stability as one of their guiding principles to facilitate communication, by conserving well-established taxonomic names in their conventional meaning (ICZN, 1999, Preamble). Mayr and Ashlock (1991) stressed the role of taxonomic stability in information retrieval because taxon names store ample information about members of the group such as their diagnostic traits, ecological niche, and distribution. New taxon names are commonly proposed as phylogenetic studies pervade the scientific literature (e.g., Zhang et al., 2020). Although establishing a new taxonomic name may be attractive, authors should explicitly acknowledge the weighting that they attribute to the stability criterion when they propose new taxonomic groups, since changes in existing names can bring considerable disruption to established research fields.

– *Morphological support* refers to the phenotypic recognizability of a group. This criterion is connected to the distinctiveness criterion but refers more specifically to the presence of diagnostic phenotypic traits, or synapomorphies, defining a group. The criterion serves as an additional line of evidence for the ‘naturalness’ of a taxonomic group, since related organisms are expected to share derived traits. This criterion depends on the perceptual modalities of humans (e.g., sight more than odor) and is important because humans are the users of the taxonomic names. Mayr and Ashlock (1991) included this measure in their degree of difference and grade characteristics criteria, and warned that convergent traits can be misleading for establishing evolutionary classifications. Vences et al. (2013) listed phenotypic recognizability as one of their three primary criteria for naming taxa, and argued that diagnostic morphological traits should ideally be externally observable and identifiable by non-specialists.

– *Clade age* criterion is based on the assumption that taxonomic groups take a certain amount of time to emerge and establish. This is a relatively recent criterion that has become more popular with the introduction of time-calibrated phylogenetic analyses. It is a seemingly straightforward criterion that Vences et al. (2013) refer to as “time banding” because it involves using a “band of time” as a cut-off value for delimiting taxonomic groups. While comparability of evolutionary age across taxonomic groups is an obvious advantage of this criterion, it is important to keep in mind that divergence times are highly labile values that can vary greatly between studies depending on the method of phylogenetic inference, analysis settings like calibration points, taxonomic sampling, and type of data used to reconstruct the phylogeny (Avise and Liu, 2011, Kraichak et al., 2017). Use of a strict time band implies uniformity of analyses between studies, as well as of detectability of evolutionary divergence rate variation among clades.

– *Distribution* criterion refers to the geographical distribution of a taxonomic group, or the restrictedness/uniqueness of their geographic range. The importance of this criterion is usually higher for lower taxonomic ranks (species, genus) and becomes less important as taxa become larger and more inclusive. Vences et al. (2013) considered biogeography to be of limited practical value due to the arbitrary nature of this criterion; however, biogeography is widely used to establish classifications for conservation and management purposes.

– *Ecological similarity* refers to distinctness of the adaptive zone or niche occupied by members of a taxonomic group. It may overlap with other criteria such as phenotypic recognizability when morphological traits evolve in response to shifts to new adaptive zones. With ecological factors being increasingly recognized as key factors in speciation and diversification of taxa (Nosil, 2012), this criterion is likely to become more important for establishing taxonomic classifications in new systematic studies.

### Criteria weights

TaxonomR assigns weights to each criterion individually on a scale of 0 to 1, depending on the importance of any given criterion. A weight of 0 indicates that a criterion has no influence on a taxonomic ranking decision and a weight of 1 is assigned to a criterion of a primary importance. Individual scores are calculated for each criterion based on how well a taxon fulfills that criterion. Details about individual score calculations are provided in Supplementary data. The final score for each taxonomic group is calculated as the fraction of the sum of scores that a group receives and the maximum possible sum of scores under a specified weighting scheme. If several alternative taxonomic groups are provided in the data file, classification combinations are scored based on the average score received by each individual group included in that combination.

### Case study: genus-level classification of fritillary butterflies

We use a phylogeny with estimated divergence times from De Moya et al. (2017) that was generated from one mitochondrial and four nuclear genes (a total of 4,307 bp) for 56 ingroup taxa (36 distinct species, 52 taxa considering subspecies) and seven outgroup taxa from the genera *Brenthis* and *Issoria*. This phylogeny was chosen because of its comprehensive sampling for both Eurasian and North American species; we expect that newer phylogenetic studies with extensive and robust molecular sequencing will provide alternate phylogenies (e.g. Campbell et al. (2020) for North American species). We started our evaluation with a “flat” weighting scheme under which all criteria are considered equally important in making a taxonomic ranking decision, with a weighting of 0.1. We set the clade age range to 8-10 MYA, which has been recently discussed as a threshold for genus age (Zhang et al., 2019). We then considered four theoretical taxonomic schemes, stressing: A) monophyly + branch length + node support (i.e., phylogenetic focus), B) usage + distribution (historical taxonomic inertia), C) usage + morphology (traditional phenetic focus), and D) clade age (molecular dating focus). For each of these we iteratively increased weights of the pertinent criteria to 0.2, 0.4, 0.6, 0.8, and 1.0 and recorded the magnitude and direction of change of the score of each taxonomic arrangement (Figure 2). Specific taxon data (clade members, stability (usage), clade age, etc.) are provided for each criterion in Table 1 and the tree and data files uploaded to TaxonomR are provided in supplemental files 1 and 2.

**Figure 2.**
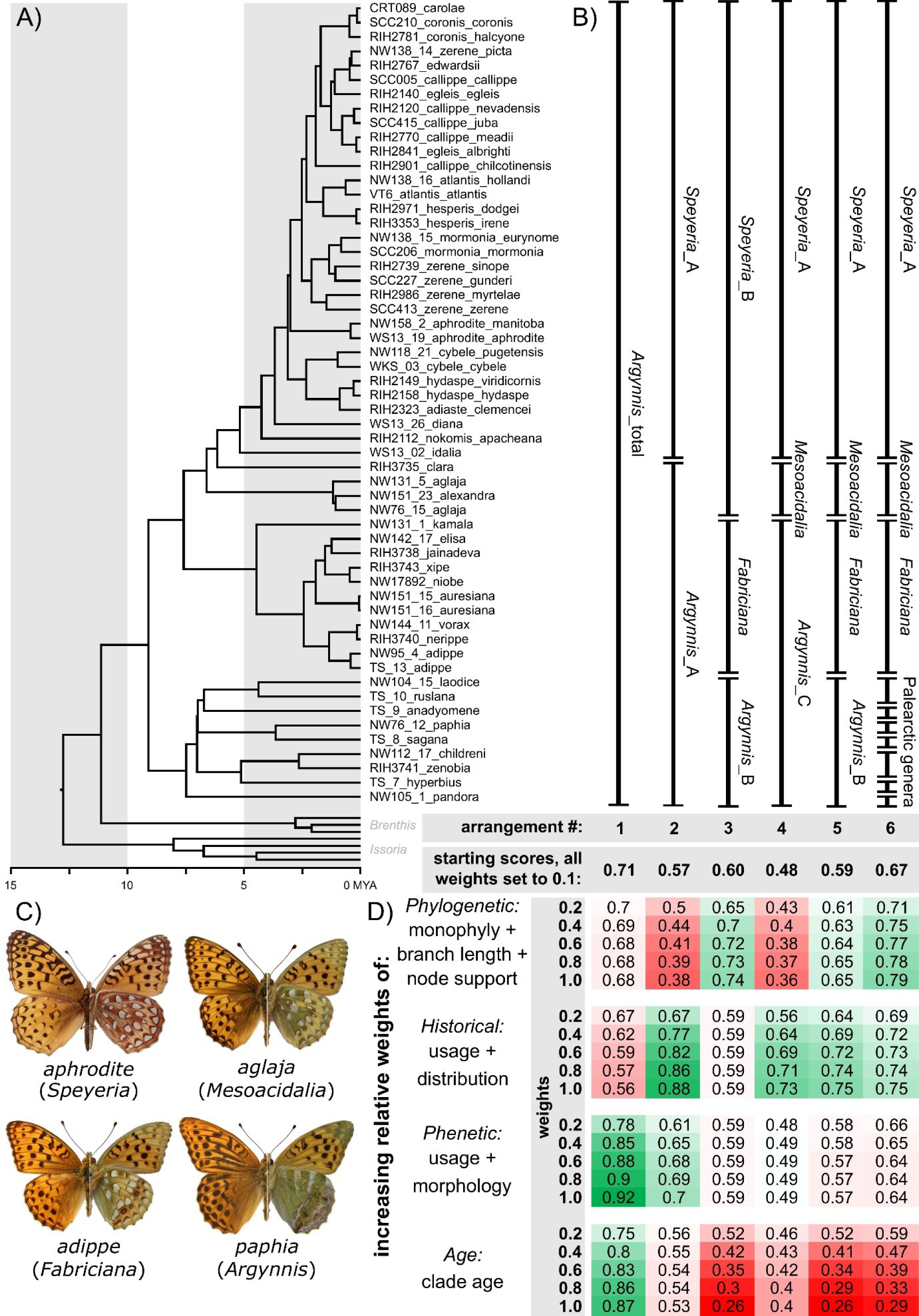
Summarized results of the greater fritillary case study. A) dated phylogeny from De Moya *et al*. (2017), B) competing taxonomic arrangements considered here, C) representative habitus images (dorsal on the left, ventral on the right) of the main groupings of greater fritillaries (for each, species epithet is above and least inclusive genus below), and D) heatmap showing resulting combined scores for each of the competing taxonomic arrangements (as shown in B) given up-weighting of specific criteria. In D, green and red indicate an increase or decrease in score, respectively, relative to the flat weighting of scores (“starting scores, all weights set to 0.1”). Photographs used with permission from Thomas Simonsen and JRD.

**Table 1.** Alternate taxonomic groups and summary of characteristics used in case study (from **tree file** and **data file**), including short description of taxa in each group, phylogenetic characteristics (monophyly, node support, branch length), stability criteria (current usage), phenotypic recognizability (morphology), clade age, and geographic distribution. Columns with † are included in the data file for this case study. Morphology is based on analysis of Simonsen 2006 and descriptions of Gorbunov (2001).

| Group | Taxa/Description | Monophyly<br>† | Node<br>support* | Branch<br>length** | Current<br>usage† | Morphology<br>† | Clade age,<br>MYA† | Distribution† |
| --- | --- | --- | --- | --- | --- | --- | --- | --- |
| *Argynnis*_total | all species | yes | 1 | 0 | major_use | synap | 9.104 | North_America, Eurasia |
| *Argynnis*_A | all Palearctic species | no | 0 | 0 | major_use | none | 9.104 | Eurasia |
| *Argynnis*_B | excludes *Fabriciana* & *Speyeria*_B | yes | 1 | 0 | minor_use | none | 7.482 | Eurasia |
| *Argynnis*_C | excl. only *Speyeria*_B | no | 0 | 0 | minor_use | none | 9.104 | Eurasia |
| *Fabriciana* | Palearctic subgroup | yes | 1 | 1 | major_use | synap | 4.467 | Eurasia |
| *Mesoacidalia* | Palearctic subgroup | no | 0 | 0 | minor_use | none | 6.596 | Eurasia |
| *Speyeria*_A | only North America | yes | 1 | 0 | major_use | synap | 5.177 | North_America |
| *Speyeria*_B | includes *Mesoacidalia* | yes | 1 | 0 | minor_use | synap | 6.596 | North_America, Eurasia |
| PAL‡ *Argynnis* | *paphia* | NA | 1 | 1 | minor_use | none | 3.646 | Eurasia |
| PAL‡ *Argyreus* | *hyperbius* | NA | 1 | 1 | minor_use | synap | 5.129 | Eurasia |
| PAL‡ *Argyronome* | *laodice*, *ruslana* | yes | 1 | 0 | minor_use | none | 4.383 | Eurasia |
| PAL‡ *Childrenia* | *childreni*, *zenobia* | yes | 1 | 0 | minor_use | synap | 2.644 | Eurasia |
| PAL‡ *Damora* | *sagana* | NA | 1 | 1 | minor_use | synap | 3.646 | Eurasia |
| PAL‡ *Nephargynnis* | *anadyomene* | NA | 1 | 1 | minor_use | synap | 6.732 | Eurasia |
| PAL‡ *Pandoriana* | *pandora* | NA | 1 | 1 | minor_use | synap | 7.482 | Eurasia |

\*Node support – posterior probability values from time-calibrated phylogeny of De Moya et al. 2017 used in this study (values rounded to 2 significant digits).
\*\*Branch supporting the clade is longer than the longest branch within that clade (intraclade divergence is lower than interclade divergence).
‡PAL refers to extensive splitting of genera in Eurasia, as discussed in the main text and labeled as “Palearctic genera” in Figure 2. Sampling by De Moya et al. 2017 included only single species of some genera in Figure 2. Morphology for PAL clades follows genitalia characterizations by Gorbunov (2001).

Given long-standing contention over taxonomic classification of *Argynnis* s.l., we focus on several widely used taxonomic arrangements within the phylogenetic context of De Moya *et al*.’s (2017) phylogeny for the Argynnini (Figure 2, top left). Arrangement #1 (*Argynnis*_total = Ar1) was the standard prior to the 1940’s (see below), and has been supported more recently by Simonsen (2006) and Zhang et al. (2020), and implemented in the (Wikipedia, 2023b) page for “*Argynnis*”. Arrangement #2 (Speyeria_A + Argynnis_A = Ar2) represents the widespread taxonomic treatment of *Speyeria* as distinct in North America following the publication of Dos Passos and Grey (1945), combined with the concurrent practice of many regional works in Eurasia listing all species in that continent as belonging to one genus (e.g., Tuzov, 2000).

Arrangement #3 (*Speyeria*_B + *Fabriciana* + *Argynnis*_B = Ar3) corresponds to the preferred configuration of De Moya et al. (2017), which is now also followed by Savela (2023), and the Wikipedia (2023a) listing for Argynnini, as well as numerous current lists of butterflies available on Wikipedia (e.g., “List of butterflies of Russia”). Arrangements #4 and #5 have not been used historically but were chosen to show the performance of TaxonomR when *Mesoacidalia* is recognized within arrangements that are otherwise similar to #2 and #3. Arrangement #6 (Ar6) shows a global compilation of the multiple genera that were in widespread regional use in Eurasia between the 1940’s and the 2000’s: e.g., Smart (1975).

## Results

An example screen shot of the TaxonomR shiny app running in a web browser is provided in Figure 1. The leftmost panel contains file uploads and slider bars to increase/decrease relative weighting of particular characteristics. In the rightmost panel, the overall phylogeny, a subtree showing a selected group, and individual group and combination group scores are displayed, and are actively updated if slider bars are adjusted.

### Fritillary butterfly case study

With De Moya *et al*.’s (2017) phylogeny for *Argynnis* s.l., we used the flat-weighting scheme (all scores set to 0.1) as a reference for comparing alternative weightings, and with this scheme, the total group *Argynnis* comprising all ingroup taxa (Ar1) was supported as the preferred classification with a score of 0.71, followed by a score of 0.67 for Ar6 (Figure 2, “starting scores, all weights set to 0.1”).

In our manipulations of criterion weightings, Ar1 consistently scored substantially higher than alternative classifications when traditional phenetic criteria (usage + morphology) as well as molecular dating (clade age) were prioritized over other criteria. Similarly, upweighting monophyly and node support criteria (data not shown), favoured Ar1. However, due to the relatively high intra-clade divergence within the total group *Argynnis*, the branch length criterion was scored 0 for noncompliance and Ar1 was not supported as the best classification under the phylogenetic framework (monophyly + node support + branch length). Instead, alternative arrangements comprising smaller phylogenetically distinct groups (Ar3, Ar6) that had scores of 1 for branch length gave higher scoring classifications. Of these, Ar6 includes a non-monophyletic group (*Mesoacedalia*) but had more single-taxon groups with the branch length criterion scored as “1”, which gave a higher total score compared to that of Ar3, which was comprised of three well-supported monophyletic groups (*Argynnis*_A + *Fabriciana* + *Speyeria*_B), but only one group (*Fabriciana*) having a maximum support for the branch length criterion.

The Ar2 arrangement has been widely used in concurrent regional classifications that recognized all species in Eurasia as *Argynnis* and all in North America as *Speyeria.* Ar2 includes a non-monophyletic *Argynnis*_A, and this classification scored poorly under the flat weighting scheme (2^nd^ lowest score), with even lower support under a phylogenetic taxonomic approach with monophyly-related criteria upweighted. Time-banding taxonomy was also less favourable for this arrangement because the group age of *Speyeria*_A falls outside of the specified time band. The strongest support for Ar2 was provided by the distribution criterion emphasizing the importance of disjunct distribution between *Speyeria* and *Argynnis* (usage + distribution). However, Ar2 only scored better than other arrangements when the usage + distribution criteria were weighted 4X higher than all other criteria (i.e., only when this combination of criteria was upweighted to 0.4, as compared to the flat scoring of 0.1, was the score for Ar2 markedly higher (0.77) than other arrangements (Figure 2)).

When all criteria were weighted equally at 0.1, the Ar3 taxonomic arrangement, which is currently widely used and recommended by De Moya et al. (2017), had a higher score than that of Ar5, which includes nonmonophyletic *Mesoacidalia*. Ar3 had an even better score when greater weight was placed on the combination of monophyly + branch length + node support criteria. The inclusion of two nonmonophyletic taxa in Ar4 (*Mesoacidalia* and *Argynnis*_C) also reduced the overall score for this arrangement substantially. However, upweighting usage + distribution gave Ar4, Ar5, and Ar6 better scores than Ar3, since *Mesoacidalia* has been more widely used in older classifications.

Finally, Ar6 is a multi-group classification in which most of the proposed genus-level groups are monophyletic and this arrangement was well-supported under the phylogenetic taxonomic approach which prioritizes monophyly and genetic distinctiveness (monophyly + node support + branch length). However, due to a greater number of low group ages and non-unique morphological traits among the groups, Ar6, as well as Ar3, Ar4, and Ar5, arrangements had low support under morphology-based taxonomic weighting and particularly under time banding.

Taken together, if monophyly and genetic distinctiveness are to be prioritized in the taxonomic decision making, Ar3 and Ar6 are the best supported genus-level classifications. Emphasizing disjunction of geographical ranges and current usage of taxonomic names favours Ar2. Upweighting either morphological distinctiveness and stability (usage) or clade age gives more support to Ar1. The time-banding approach to taxonomic decisions appears to be the most volatile and difficult to assess because it depends on multiple factors (e.g., mean estimate of the divergence ages, width of the time band) that vary substantially depending on the data type and methodology used to infer divergence dates, as well as relatively arbitrary prior decisions on ideal clade ages.

## Discussion

As a field, taxonomy embodies an enigmatic mix of objectivity and subjectivity. On the one hand, personal taxonomic expertise may require years of time commitment to develop, with the delimitation and ranking of named groups becoming an exercise where a taxon is whatever a taxonomist says it is. On the other hand, many argue for more objective criteria to be used in taxonomic revisions (Condamine et al., 2023, Mayr and Ashlock, 1991, Vences et al., 2013) and may even advocate for focussed reliance on single criteria like clade age (Zhang et al., 2019). Into this fray, we present TaxonomR, an interactive, web-hosted decision support tool to evaluate and quantify alternative taxonomic arrangements. The utility of TaxonomR lies in its consideration of criteria commonly used in taxonomic science, but in a quantitative manner where different criteria can be given relative weights reflecting their importance to the scoring of various taxonomic groupings. The interactive nature of TaxonomR allows both detailed exploration of a group’s alternative taxonomic arrangements and improves reproducibility and replicability of a field that is often thought of as inconsistent and subjective. Additionally, the interactive nature of TaxonomR provides a tool for teaching taxonomic and systematic principles to students and taxonomists in training, which we hope will contribute toward ameliorating the taxonomic impediment (Engel et al., 2021). In this context, it is important to consider logistical considerations, constraints, and strengths of TaxonomR as well as how this tool could advance objectivity and communication in taxonomy in a broad sense.

Given the inconsistency of taxonomic ranks (e.g., Avise and Liu, 2011, Kraichak et al., 2017), the utility of TaxonomR intrinsically depends on the complexity of a given group’s taxonomic history and characteristics of their evolution. Our case study with fritillary butterflies represents a moderately complex case of taxonomic instability that allowed us to demonstrate several of the uses of this tool. But some considerations are important to reiterate about this tool’s utility. First, upweighting phylogenetic and clade age criteria are completely reliant on the phylogeny in question and the relative sizes and ages of particular clades. For example, in our case study, the paraphyly of *Mesoacidalia* had a less pronounced effect on combined group scores than the larger and older paraphyletic clade of *Argynnis*_A; likewise, our use of a clade age of 8-10 MYA optimized scoring of clades falling within that age range given this specific dated tree. This age range has been recently posed in studies of various butterfly genera, including discussions of *Argynnis*/*Speyeria* (De Moya et al., 2017, Zhang et al., 2019), which was our rationale for selecting this age range. However even this rationalized choice affected our results and may be an impetus to test several age range bands. For example, our *Argynnis*_B clade falls just shy of this threshold (7.482 MYA, Table 1), as opposed to *Childrenia* which was far below the threshold (2.644 MYA), yet these two clades were scored identically for clade age in our treatment. Alternative clade age ranges from molecular dating studies are commonly of interest in studying higher-level group phylogenies, and TaxonomR’s interactive slider bars make it feasible to consider alternative dating prior thresholds for various genus-ranked groups (e.g., in Nymphalidae: Wahlberg (2006), Wahlberg *et al*.(2009); Papilionidae: Condamine *et al*. 2018, 2023; Squamates: Simoes *et al*. (2020); cichlids: Matschiner et al. (2020); ray-finned and bony fishes: Hughes et al. (2018), Betancur-R et al. (2017)). Most ecological characteristics, such as host plant/habitat use, are generally uniform or equally variable across our focal group of fritillary butterflies, so we did not consider any ecological characteristics here. However, TaxonomR provides substantial expanded functionality that may be applied to groups with more diverse ecologies or morphologies (e.g., ray-finned fishes, Hughes et al. (2018); Clupeiform fishes, Wang et al. (2022)).

We developed TaxonomR to provide additional objectivity to the field of taxonomy, particularly for weighting and comparing competing higher-level taxonomic classifications (e.g., Condamine et al., 2023, Zhang et al., 2019). For example, expected clade age and prior usage + morphology supported recognition of all species in a single genus (*Argynnis*), while the three-genus arrangement of De Moya et al. (2017) was only slightly favoured when monophyly, branch length and node support were emphasized. These results depend on the input file, and recognition of that dependency is a point where a structured decision making process can be initiated to generate and validate such an input file by a group of interested taxonomists, thereby fostering communication, cooperation, and community as part of structured decision making.

Additionally, the approach that underlies TaxonomR presents a unique opportunity to assess historical taxonomic classifications in a framework that allows a straightforward comparison and evaluation of how prioritization schemes might have changed over time. Just as TaxonomR can be used going forward to provide supporting justification for one classification arrangement over another, this tool can just as easily be used to go back in time to infer what criteria and lines of reasoning are likely to have been emphasized in historically favoured taxonomic arrangements. Creating such a retrospective connection in taxonomy is essential for preserving centuries worth of natural history research legacy.

Increased replicability and understanding of the processes that contribute to it will facilitate communication among taxonomists as well as with the end-users of taxonomy, and ultimately increase the democratization of taxonomy by outlining the base assumptions and criteria used by taxonomists. Ideally, this tool will allow us to view taxonomy as a Bayesian process, where taxonomic decisions represent iterative adjustments to how individual taxonomists and the broader community treat particular species or groups. By quantifying these iterative adjustments, those taxonomic changes are grounded to a commonality that allows taxonomists to acknowledge uncertainty and potential taxonomic instability. Instead of unilaterally breaking a clade or grouping based on a particular biological or phylogenetic characteristic, future taxonomic studies can present alternative arrangements and their quantifications, thus conceptually addressing the confidence intervals of alternative taxonomies.

## Acknowledgements

We thank Thomas Simonsen for photographs of Palearctic representatives used in Figure 2. This research was supported by an NSERC Discovery Grant to FAHS (#RGPIN-2018-04920) and USDA-NIFA HATCH grant to JRD (Project KY008091).

## Author contributions

All authors conceptualized software and contributed to the manuscript writing and editing. OVV implemented software with input from JRD and FAHS.

## Data availability

Software, documentation, and example data are available at https://github.com/OksanaVe/TaxonomR and as an online application at https://oksanav.shinyapps.io/TaxonomR/.

## References

Agnarsson, I. & Kuntner, M. 2007. Taxonomy in a changing world: seeking solutions for a science in crisis. Systematic biology, 56, 531–539.

Allison, D. B., Shiffrin, R. M. & Stodden, V. 2018. Reproducibility of research: Issues and proposed remedies. Proceedings of the National Academy of Sciences, 115, 2561–2562.

Avise, J. C. & Liu, J.-X. 2011. On the temporal inconsistencies of Linnean taxonomic ranks. Biological Journal of the Linnean Society, 102, 707–714.

Baker, M. 2016. Reproducibility crisis. Nature, 533, 353–66.

Bessette, D., Wilson, R., Beaudrie, C. & Schroeder, C. 2019. An online decision support tool to evaluate ecological weed management strategies. Weed Science, 67, 463–473.

Betancur-R, R., Wiley, E. O., Arratia, G.,Acero, A., Bailly, N., Miya, M., Lecointre, G. & Orti, G. 2017. Phylogenetic classification of bony fishes. BMC evolutionary biology, 17, 1–40.

Bhargava, H. K., Power, D. J. & Sun, D. 2007. Progress in Web-based decision support technologies. Decision Support Systems, 43, 1083–1095.

Braby, M. F., Eastwood, R. & Murray, N. 2012. The subspecies concept in butterflies: has its application in taxonomy and conservation biology outlived its usefulness? Biological Journal of the Linnean Society, 106, 699–716.

Campbell, E. O., Gage, E. V., Gage, R. V. & Sperling, F. A. H. 2020. Single nucleotide polymorphism-based species phylogeny of greater fritillary butterflies (Lepidoptera: Nymphalidae:Speyeria) demonstrates widespread mitonuclear discordance. Systematic Entomology, 45, 269–280.

Condamine, F. L., Allio, R., Reboud, E. L., Dupuis, J. R., Toussaint, E. F., Mazet, N., Hu, S.-J., Lewis, D. S., Kunte, K. & Cotton, A. M. 2023. A comprehensive phylogeny and revised taxonomy illuminate the origin and diversification of the global radiation of Papilio (Lepidoptera: Papilionidae). Molecular Phylogenetics and Evolution, 183, 107758.

Condamine, F. L., Nabholz, B., Clamens, A. L., Dupuis, J. R. & Sperling, F. A. 2018. Mitochondrial phylogenomics, the origin of swallowtail butterflies, and the impact of the number of clocks in Bayesian molecular dating. Systematic entomology, 43, 460–480.

Copp, G. H., Vilizzi, L., Wei, H., Li, S., Piria, M., Al-Faisal, A. J., Almeida, D., Atique, U., Al-Wazzan, Z. & Bakiu, R. 2021. Speaking their language–development of a multilingual decision-support tool for communicating invasive species risks to decision makers and stakeholders. Environmental Modelling & Software, 135, 104900.

De Moya, R. S., Savage, W. K., Tenney, C., Bao, X., Wahlberg, N. & Hill, R. I. 2017. Interrelationships and diversification of A rgynnis F abricius and S peyeria S cudder butterflies. Systematic Entomology, 42, 635–649.

Dos Passos, C. F. & Grey, L. P. 1945. A genitalic survey of Argynninae (Lepidoptera, Nymphalidae). American Museum novitates; no. 1296.

Engel, M. S., Ceríaco, L. M., Daniel, G. M., Dellapé, P. M., Löbl, I., Marinov, M., Reis, R. E., Young, M. T., Dubois, A. & Agarwal, I. 2021. The taxonomic impediment: a shortage of taxonomists, not the lack of technical approaches. Oxford University Press UK.

Garnett, S. T. & Christidis, L. 2017. Taxonomy anarchy hampers conservation. Nature, 546, 25–27.

Garnett, S. T. & Christidis, L. 2018. Science-based taxonomy still needs better governance: Response to Thomson et al. PLoS Biology, 16, e2005249.

Gorbunov, P. & Kosterin, O. 2007. The butterflies (Hesperioidea and Papilionoidea) of North Asia (Asian part of Russia) in nature, Rodina & Fodio, Aidis Producers House.

Gorbunov, P. Y. 2001. The butterflies of Russia: classification, genitelia, keys for identification, Thesis.

Hemming, V., Camaclang, A. E., Adams, M. S., Burgman, M., Carbeck, K., Carwardine, J., Chadès, I., Chalifour, L., Converse, S. J. & Davidson, L. N. 2022. An introduction to decision science for conservation. Conservation biology, 36, e13868.

Higgins, L. & Riley, N. 1983. Field Guide to the Butterflies of Britain nad Europe. London. HIGGINS, L. G. 1975. Classification of European butterflies, Collins.

Hughes, L. C., Ortí, G., Huang, Y., Sun, Y., Baldwin, C. C., Thompson, A. W., Arcila, D., Betancur-R, R., Li, C. & Becker, L. 2018. Comprehensive phylogeny of ray-finned fishes (Actinopterygii) based on transcriptomic and genomic data. Proceedings of the National Academy of Sciences, 115, 6249–6254.

ICZN 1999. International Code of Zoological Nomenclature, London, The International Trust for Zoological Nomenclature.

Kehimkar, I. D. 2008. Book of Indian butterflies, Oxford University Press.

Kraichak, E., Crespo, A., Divakar, P. K., Leavitt, S. D. & Lumbsch, H. T. 2017. A temporal banding approach for consistent taxonomic ranking above the species level. Scientific Reports, 7, 1–7.

Linnaeus, C. 1735. Systema naturae; sive, Regna tria naturae: systematice proposita per classes, ordines, genera & species, Haak.

Matschiner, M., Böhne, A., Ronco, F. & Salzburger, W. 2020. The genomic timeline of cichlid fish diversification across continents. Nature communications, 11, 5895.

Mayr, E. & Ashlock, P. 1991. Principles of systematic zoology. 2da edition. MacGraw-Hill. Inc. New York.

Morales-Torres, A., Escuder-Bueno, I., Andrés-Doménech, I. & Perales-Momparler, S. 2016. Decision Support Tool for energy-efficient, sustainable and integrated urban stormwater management. Environmental Modelling & Software, 84, 518-528.

Moritz, C. 1994. Defining ‘evolutionarily significant units’ for conservation. Trends in ecology & evolution, 9, 373–375.

Nosil, P. 2012. Ecological speciation, Oxford University Press.

Padial, J. M., Miralles, A., De La Riva, I. & Vences, M. 2010. The integrative future of taxonomy. Frontiers in zoology, 7, 1–14.

Pelham, J. P. 2023. A catalogue of the butterflies of the United States and Canada. [Online]. Available: https://www.butterfliesofamerica.com/US-Can-Cat.htm [Accessed 28 May 2023].

Power, D. J. 2007. A brief history of decision support systems. DSSResources.com, 3.

Raven, P. H., Berlin, B. & Breedlove, D. E. 1971. The Origins of Taxonomy: A review of its historical development shows why taxonomy is unable to do what we expect of it. Science, 174, 1210–1213.

Reuss, F. A. T. 1926. Systematischer Überblick der Dryadinae T. Rss. Mit einigen Neubeschribungen (Lep. Rhopal.). Deutsche Entomologische Zeitschrift 1926, 65–70.

Savela, M. 2023. Lepidoptera and other life forms. Heliconiinae. [Online]. Available: https://www.funet.fi/pub/sci/bio/life/insecta/lepidoptera/ditrysia/papilionoidea/nymphalidae/heliconiinae/ [Accessed 7 May 2023].

Schram, F. R. 2004. The truly new systematics–megascience in the information age. Hydrobiologia, 519, 1–7.

Shiffrin, R. M., Börner, K. & Stigler, S. M. 2018. Scientific progress despite irreproducibility: A seeming paradox. Proceedings of the National Academy of Sciences, 115, 2632–2639.

Shou, J., Chou, I. & Li, Y. 2006. Systematic butterfly names of the world.

Simões, T. R., Caldwell, M. W. & Pierce, S. E. 2020. Sphenodontian phylogeny and the impact of model choice in Bayesian morphological clock estimates of divergence times and evolutionary rates. BMC biology, 18, 1–30.

Simonsen, T., Wahlberg, N., Brower, A. & De Jong, R. 2006. Morphology, molecules and fritillaries: approaching a stable phylogeny for Argynnini (Lepidoptera: Nymphalidae). Insect Systematics & Evolution, 37, 405–418.

Simonsen, T. J. 2006. Fritillary phylogeny, classification, and larval host plants: reconstructed mainly on the basis of male and female genitalic morphology (Lepidoptera: Nymphalidae: Argynnini). Biological Journal of the Linnean Society, 89, 627–673.

Smart, P. 1975. Encyclopedia of the Butterfly World, New York Salamander Books.

Thomson, S. A., Pyle, R. L., Ahyong, S. T., Alonso-ZARAZAGA, M., Ammirati, J., Araya, J. F., Ascher, J. S., Audisio, T. L., Azevedo-SANTOS, V. M. & Bailly, N. 2018. Taxonomy based on science is necessary for global conservation. PLoS biology, 16, e2005075.

Tuzov, V. K. 2000. Genus Argynnis. In: TUZOV, V. K. (ed.) Guide to the Butterflies of Russia and Adjacent Territories. Sofia, Bulgaria: Pensoft.

Vences, M., Guayasamin, J. M., Miralles, A. & De La Riva, I. 2013. To name or not to name: criteria to promote economy of change in supraspecific Linnean classification schemes.

Wahlberg, N. 2006. That awkward age for butterflies: insights from the age of the butterfly subfamily Nymphalinae (Lepidoptera: Nymphalidae). Systematic Biology, 55, 703–714.

Wahlberg, N. 2020. Nymphalidae.net. Argynnini. updated 2020-11-20 [Online] [Online]. Available: http://www.nymphalidae.net/Nymphalidae/Classification/Hel_Argynnini.htm [Accessed 28 May 2023].

Wahlberg, N., Leneveu, J., Kodandaramaiah, U., Peña, C., Nylin, S., Freitas, A. V. & Brower, A. V. 2009. Nymphalid butterflies diversify following near demise at the Cretaceous/Tertiary boundary. Proceedings of the Royal Society B: Biological Sciences, 276, 4295–4302.

Wang, Q., Dizaj, L. P., Huang, J., Sarker, K. K., Kevrekidis, C., Reichenbacher, B., Esmaeili, H. R., Straube, N., Moritz, T. & Li, C. 2022. Molecular phylogenetics of the Clupeiformes based on exon-capture data and a new classification of the order. Molecular Phylogenetics and Evolution, 175, 107590.

Warren, A. D., Davis, K. J., Stangeland, E. M., Pelham, J. P., Willmott, K. R. & Grishin, N. V. 2023. Illustrated Lists of American Butterflies [Online]. Available: http://www.butterfliesofamerica.com/US-Can-Cat.htm [Accessed 7 May 2023].

Warren, B. 1955. A review of the classification of the subfamily Argynninae (Lepidoptera: Nymphalidae). Part 2. Definition of the Asiatic genera. Transactions of the Royal Entomological Society of London, 107, 381-392.

Warren, B. C. 1944. Review of the classification of the Argynnidi: with a systematic revision of the genus Boloria. Trans. Royal ent. Soc. Lond., 94, 1–101.

Wheeler, Q. D. 2008. The new taxonomy, CRC Press.

Wheeler, Q. D., Raven, P. H. & Wilson, E. O. 2004. Taxonomy: impediment or expedient?: American Association for the Advancement of Science.

WIKIPEDIA. 2023a. Argynnini [Online] [Online]. Available: https://en.wikipedia.org/wiki/Argynnini [Accessed 7 May 2023].

WIKIPEDIA. 2023b. Argynnis [Online] [Online]. Available: https://en.wikipedia.org/wiki/Argynnis [Accessed 28 May 2023].

Wilby, R. L., Dawson, C. W. & Barrow, E. M. 2002. SDSM—a decision support tool for the assessment of regional climate change impacts. Environmental Modelling & Software, 17, 145–157.

Wilson, E. O. 2004. Taxonomy as a fundamental discipline. Philosophical Transactions of the Royal Society of London. Series B: Biological Sciences, 359, 739–739.

Zhang, J., Cong, Q., Shen, J., Brockmann, E. & Grishin, N. V. 2019. Genomes reveal drastic and recurrent phenotypic divergence in firetip skipper butterflies (Hesperiidae: Pyrrhopyginae). Proceedings of the Royal Society B, 286, 20190609.

Zhang, J., Cong, Q., Shen, J., Opler, P. A. & Grishin, N. V. 2020. Genomic evidence suggests further changes of butterfly names. The taxonomic report of the International Lepidoptera Survey, 8.

