## Supplemental data for "Toward transparent taxonomy: an interactive web-tool for evaluating competing taxonomic arrangements"

*Criteria weights*

TaxonomR assigns weights to each criterion individually on a scale of 0 to 1, depending on the importance of any given criterion. A weight of 0 indicates that a criterion has no influence on a taxonomic ranking decision and a weight of 1 is assigned to a criterion of a primary importance. Individual scores are calculated for each criterion based on how well a taxon fulfills that criterion. Details about :

- Monophyly: if a taxonomic group is monophyletic, a taxon receives a full weight score; if a group is not monophyletic the score for this criterion is set to 0;
- Node support score is a continuous variable that is proportional to the node support value converted to a decimal value between 0 and 1. The score is calculated as product of weight and clade support values, e.g., a clade with bootstrap support of 80% and criterion weight set to 1 (primary importance) has a criterion score of 0.8 x 1 = 0.8. If a group is not monophyletic, the clade support score is 0.
- Branch length score is calculated as a ratio between the longest internal branch within a clade and that of the stem branch leading to the group. If the ratio is greater than one (intra-clade divergence is greater than divergence between clades), the score is 0; if the ratio equals one (equal degree of intra- and inter-clade divergence), the score is 0.5; if the ratio is less than one, the score is 1. For single-taxon groups, the branch length score is scored as 1.
- Current usage (stability) score is calculated based on which of three discrete categories is assigned to a taxonomic group. If a group is scored as “major_use”, the stability score equals to the full weight value, a “minor_use” category receives 50% of the user-defined weight, “new_use” category receives a score of 25% of the weight value, and “new” category receives a stability score of 0.
- Morphological support score is determined based on which of the four categories a group is assigned to. This score equals 100, 50, 25, or 0 per cent of the weight value for the “synap”, “combo”, “plastic”, and “none” categories, respectively.
- Clade age score equals that of the full weight value if the age of a taxonomic group is within a user defined time band. If the age of a clade falls outside the specified time range, this criterion score equals 0. In its current implementation, TaxonomR does not assign partial weights to groups that fall outside of the user-specified age band regardless of how close the groups are to the time band, thus the age band for a particular user case should be considered carefully. The initial age range is set automatically with its median value equal to the specified age cut-off specified in the data file.
- Distribution score is calculated based on the restrictiveness of a biogeographical range of a taxonomic group. If distribution of a group is restricted to a single biogeographical unit, the score is equal to the full weight value; if members of a group occur in two or more biogeographical units, the score is proportional to the fraction of all specified distribution units in which a taxon occurs and is calculated as a function:

$s= \frac{(n-u)}{n}*w$,

Where *s* is a group’s biogeography score, *n* is a total number of unique biogeographical units specified across all groups in a study, *u* is the number of distribution units in which members of any particular taxon occur, and *w* is a user specified weight of the distribution criterion. For a widely distributed taxon that occurs in all distribution units, *u* = *n* and *s* = 0, indicating that a taxon does not have a restricted distribution.

- Ecological similarity and miscellaneous trait scores are calculated for each trait individually (up to three traits maximum); each trait is calculated in a manner similar to the distribution criterion based on how restricted/specialized a group is for a specified functional trait.

*Final total scores*

The final score for each taxonomic group is calculated as the fraction of the sum of scores that a group receives and the maximum possible sum of scores under a specified weighting scheme. In this way, final scores are standardized and expressed as decimal values on a scale from 0 to 1. Groups are then sorted based on their final total score and displayed in descending order. Additionally, groups can be highlighted and visually expanded by selecting a group from a drop-down menu. This feature is useful when working with large phylogenies that make it difficult to see the composition of the group and its placement in a tree.

*Group combination scores*

If several alternative taxonomic groups are provided in the data file, classification combinations are scored based on the average score received by each individual group included in that combination. Taxonomic group combinations are sorted and listed according to their scores.
